# *Tet1* isoforms differentially regulate gene expression, synaptic transmission and memory in the mammalian brain

**DOI:** 10.1101/2020.07.27.223685

**Authors:** C.B. Greer, J. Wright, J.D. Weiss, R.M. Lazerenko, S.P. Moran, J. Zhu, K.S. Chronister, A.Y. Jin, A.J. Kennedy, J.D. Sweatt, G.A. Kaas

## Abstract

The dynamic regulation of DNA methylation in post-mitotic neurons is necessary for memory formation and other adaptive behaviors. Ten-eleven translocation 1 (TET1) plays a part in these processes by oxidizing 5-methylcytosine (5mC) to 5-hydroxymethylcytosine (5hmC), thereby initiating active DNA demethylation. However, attempts to pinpoint its exact role in the nervous system have been hindered by contradictory findings, perhaps due in part, to a recent discovery that two isoforms of the *Tet1* gene are differentially expressed from early development into adulthood. Here, we demonstrate that both the shorter transcript (*Tet1*^*S*^) encoding an N-terminally truncated TET1 protein and a full-length *Tet1* (*Tet1*^*FL*^) transcript encoding canonical TET1 are co-expressed in the adult brain. We show that *Tet1*^*S*^ is the predominantly expressed isoform, and is highly enriched in neurons, whereas *Tet1*^*FL*^ is generally expressed at lower levels and more abundant in glia, suggesting their roles are at least partially cell-type specific. Using viral-mediated, isoform- and neuron-specific molecular tools, we find that *Tet1*^*S*^ repression enhances, while *Tet1*^*FL*^ impairs, hippocampal-dependent memory. In addition, the individual disruption of the two isoforms leads to contrasting changes in basal synaptic transmission and the dysregulation of unique gene ensembles in hippocampal neurons. Together, our findings demonstrate that each *Tet1* isoform serves a distinct role in the mammalian brain.

## Introduction

DNA methylation is an essential regulator of gene expression in the brain, and is required for learning and memory formation (Jarome and Lubin, 2014). Based on its role during development, DNA methylation was initially thought to function as a stable epigenetic mark in post-mitotic cells in the brain, but it is now know to be dynamically regulated—in response to neuronal stimulation, learning, and experience (Martinowich et al., 2003; Miller and Sweatt, 2007; Saunderson et al., 2016). DNA methylation levels are controlled by the antagonistic actions of DNA methyltransferases (DNMTs), which methylate the 5th carbon of cytosine bases (5mC) (Okano et al., 1999; Hermann et al., 2004), and the Ten-Eleven translocation (TETs) enzymes, which oxidize 5mCs to 5-hydroxymethylcytosine (5hmC) and initiate active DNA demethylation (Tahiliani et al., 2009; Guo et al., 2011). TET enzymes are critical for brain function and mutations or changes in the expression of *Tet* genes are associated with, or the cause of, cognitive deficits in humans (Dong et al., 2015; Cochran et al., 2019; Beck et al., 2020). Thus, the study of TET-mediated mechanisms may provide novel insights into the pathophysiology of neurological disease.

All three *Tet* genes *(Tet1-3)* are expressed in the mammalian brain and studies suggest they generally serve non-redundant functions. *Tet3* is the highest expressed and is transcriptionally upregulated by neuronal stimulation (Widagdo et al., 2014). Knockdown of *Tet3* alters synaptic transmission, and conditional knockout (KO) of the gene impairs spatial memory, indicating that its necessary for cognition (Yu et al., 2015; Antunes et al., 2020). *Tet2* is also abundantly expressed in the brain and its disruption is associated with enhanced spatial memory, suggesting it may function as a negative regulator in the brain (Zengeler et al., 2019). *Tet1*, despite its much lower expression, has been the most studied *Tet* family member in the nervous system and is implicated in the regulation of activity-dependent gene expression, synaptic transmission and cognition (Alaghband et al., 2016). However, attempts to define its exact role, particularly in the context of learning and memory, have been hampered by inconsistent findings. For instance, depending on the study, loss of the gene in KO mice has been reported to either impair, enhance, or have no effect, on memory (Rudenko et al., 2013; Zhang et al., 2013; Kumar et al., 2015; Towers et al., 2018). Likewise, overexpression of *Tet1* enhances memory, while expression of its catalytic domain does the opposite (Kaas et al., 2013; Kwon et al., 2018). A potential explanation for some of these inconsistences comes from a recent report that the *Tet1* gene undergoes an isoform switch from the full-length, canonical transcript (hereafter *Tet1*^*FL*^) in embryonic stem cells to a shorter, truncated variant (hereafter *Tet1*^*S*^) exclusive to somatic tissues (Zhang et al., 2016). In addition, evidence suggests that in some tissues both transcripts might be co-expressed (Good et al., 2017; Yosefzon et al., 2017). Whether this is the case in the adult brain, and if so, what functions these *Tet1* isoforms might serve, has not been explored.

Here we report that both *Tet1* isoforms are expressed in the brain. The *Tet1*^*S*^ isoform is highly enriched in neurons and its expression is regulated in an activity-dependent manner. In contrast, *Tet1*^*FL*^ is transcribed at low basal levels in neurons, yet expressed at much higher levels in glia. Using newly-developed molecular tools, we found that transcriptional repression of *Tet1*^*S*^ enhanced, while transcriptional repression of *Tet1*^*FL*^, impaired, long-term memory formation. Moreover, the repression of *Tet1*^*FL*^ and *Tet1*^*S*^ in neurons had opposing effects on basal synaptic transmission. Genome-wide transcriptional profiling revealed that the dissimilar effects of their disruption result, in part, from the distinct gene ensembles regulated by *Tet1*^*FL*^ and *Tet1*^*S*^. Taken together, these results strongly indicate that *Tet1*^*FL*^ and *Tet1*^*S*^ serve important, non-redundant functions in the nervous system.

## Results

### *Tet1* is expressed as two distinct transcripts in the adult brain

In order to establish whether the *Tet1* gene is expressed as more than one transcript in the adult brain, we first examined the *Tet1* 5’coding region for promoter-associated histone marks using published chromatin immunoprecipitation sequencing (ChIP-seq) datasets derived from NeuN+ hippocampal neurons (Halder et al., 2015). Two regions within the *Tet1* gene locus were enriched with H3 lysine 4 tri-methylation (H3K4me3), H3 lysine 27 acetylation (H3K27ac) and H3 lysine 9 acetylation (H3K9ac), marks typically associated with transcriptionally active promoters (Liang et al., 2004; Gates et al., 2017; Sato et al., 2019). The distal site, termed promoter 1, lied upstream of *Tet1* Refseq exon 1 and was only mildly enriched for the three epigenetic modifications, whereas the second region, termed promoter 2, was present in an intronic region just upstream of *Tet1* Refseq exon 2 and was characterized by much stronger active histone peaks (Fig. 1*A*). To test if like other active promoters, RNA polymerase II (RNAP2) was enriched at these sites, we conducted ChIP on hippocampal chromatin using primers targeted to each region. At both sites, we observed significant RNAP2 enrichment compared to the negative control region, (fold change: F (2, 15) = 5.2, p = 0.0198, One-Way ANOVA; neg. control, 1 ± 0.16 vs. site 1, 2.3 ± 0.5, p = 0.029, neg. control, 1 ± 0.16 vs. site 2, 2.4 ± 0.3, p = 0.023, Dunnett’s *post hoc;* n = 6 for all groups) indicating that each promoter was likely transcriptionally active (Fig. 1*B*).

**Figure 1.**
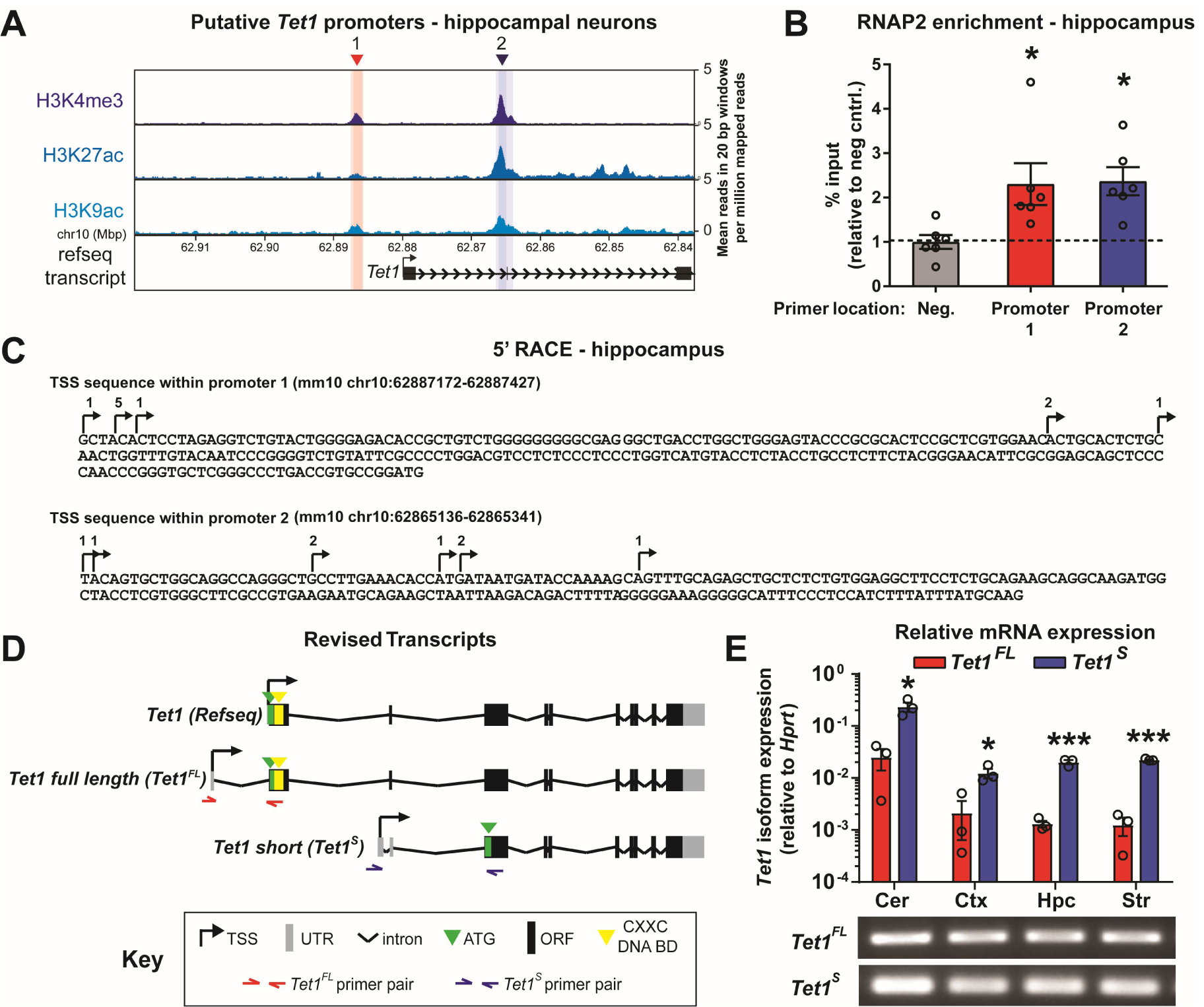
*Tet1* is expressed as two distinct transcript isoforms in the adult mouse brain. ***A***, Mean normalized H3K4me3, H3K27ac and H3K9ac ChIP-seq signal in hippocampal neurons at the *Tet1* gene locus (Halder et al., 2015). Red and blue shaded areas depict putative *Tet1* isoform promoter regions relative to the annotated RefSeq transcript. ***B***, ChIP-qRT-PCR analysis of RNAP2 enrichment (% input) at predicted *Tet1* promoters in adult hippocampal tissue relative to a negative control (mouse *Igx1a*). * p < 0.05 (Dunnett’s *post hoc*), p < 0.05 (One Way ANOVA). n = 6 mice. Data represent mean ± SEM. ***C***, 5’ RACE sequence results summary from DNA clones amplified from adult hippocampal RNA. n = 8-10 clones/isoform. Arrows and numbers above each nucleotide represent the transcriptional start sites and number of clones, respectively. ***D***, Illustration of revised *Tet1* isoform transcript architecture based on ChIP and 5’ RACE data. **Key**: grey, untranslated region (UTR); black, open reading frame (ORF); green, ATG; yellow, CXXC non-methyl CpG binding domain. Half arrows represent isoform specific primer locations. ***E***, Top: qRT-PCR analysis of *Tet1* isoform expression levels in adult brain sub-regions relative to *Hprt*. Bottom: Image of endpoint PCR products generated after qRT-PCR using *Tet1*^*FL*^ and *Tet1*^*S*^-specific primers. * p < 0.05, *** p < 0.001 (unpaired two-tailed *t* test). n = 3. Data represent mean ± SEM.

Next, we used hippocampal RNA and 5’ Rapid Amplification of cDNA Ends (RACE) to identify the transcriptional start sites (TSSs) aligned with each of the predicted *Tet1* promoters. As is common for many genes, the precise TSSs within both promoter regions varied by several nucleotides (Giardina and Lis, 1993; Leenen et al., 2015) (Fig. 1*C*). The transcript starting at promoter 1, termed *Tet1 full length (Tet1*^*FL*^), encodes the full length canonical TET1 enzyme translated from a start codon (Kozak sequence-gccATGt) located in exon 2 (Refseq *Tet1* exon 1). While the transcript arising from intronic promoter 2, termed *Tet1 short (Tet1*^*S*^), encodes for a truncated enzyme lacking a large portion of the TET1^FL^ N-terminus, including the CXXC non-methylated CpG binding domain (Fig. 1*D*). TET1^S^ is translated from a start codon (Kozak sequence-tccATGg) located in Refseq *Tet1* exon 3.

In order to examine where, and to what extent, *Tet1*^*FL*^ and *Tet1*^*S*^ were expressed in the brain, we designed isoform-specific primers and performed qRT-PCR using cDNA libraries generated from the cerebellum, cortex, hippocampus, and striatum. Both *Tet1* transcripts were detected in all four brain regions, with the mRNA levels of *Tet1*^*S*^ approximately 10-fold higher than *Tet1*^*FL*^ across all samples, indicating that its the predominant *Tet1* transcript expressed in the brain. Moreover, we found that in the cerebellum, the levels of both transcripts were an order of magnitude higher than in any other brain regions surveyed (fold change: Cer-*S* 0.23 ± 0.049, vs. Cer-*FL* 0.025 ± 0.011, t_(4)_ = 4.2, p = 0.0138; Ctx-*S*, 0.016 ± 0.0026 vs. Ctx-*FL*, 0.0021 ± 0.0015, t_(4)_ = 4.5, p = 0.0106 ; Hpc-*S*, 0.02 ± 0.0018 vs. Hpc-*FL*, 0.0013 ± 0.0002, t_(4)_ = 10, p = 0.0005; Str-*S*, 0.022 ± 0.00057 vs. Str-*FL*, 0.0013 ± 0.00049, t_(4)_ = 28, p < 0.0001; n = 3 all groups; unpaired two-tailed *t* test) (Fig. 1*E*). We also measured *Tet1*^*FL*^ and *Tet1*^*S*^ mRNA levels in the adult heart, kidney, liver, muscle, and spleen. We found both transcripts were present, and expressed at ratios comparable to those in the brain, suggesting this *Tet1* expression pattern is a general feature of most somatic tissues (data not shown). Taken together, our results demonstrate that two transcripts, encoding distinct protein isoforms, are actively generated from the *Tet1* gene in the adult mammalian brain and that the novel, truncated *Tet1*^*S*^ is the predominant transcript.

### *Tet1* isoform transcript usage significantly differs between neurons and glia

Because our initial experiments were conducted using heterogeneous brain tissue, we next compared the expression levels of *Tet1*^*FL*^ and *Tet1*^*S*^ transcripts in hippocampal neurons and glia using qRT-PCR. *Tet1*^*S*^ was expressed at ∼3-fold higher levels in neurons than in glia (fold change: *S-*glia, 0.34 ± 0.017 *vs. S-*neurons, 1 ± 0.14, p = 0.0005, n = 6; unpaired two-tailed *t* test), whereas *Tet1*^*FL*^ transcripts were ∼15-fold more abundant in glia than in neurons (fold change: *FL-*glia, 15 ± 0.74 *vs. FL-*neurons, 1.2 ± 0.32, t_(10)_ = 17, p < 0.0001, n = 6; unpaired two-tailed *t* test) (Fig. 2*A*).

**Figure 2.**
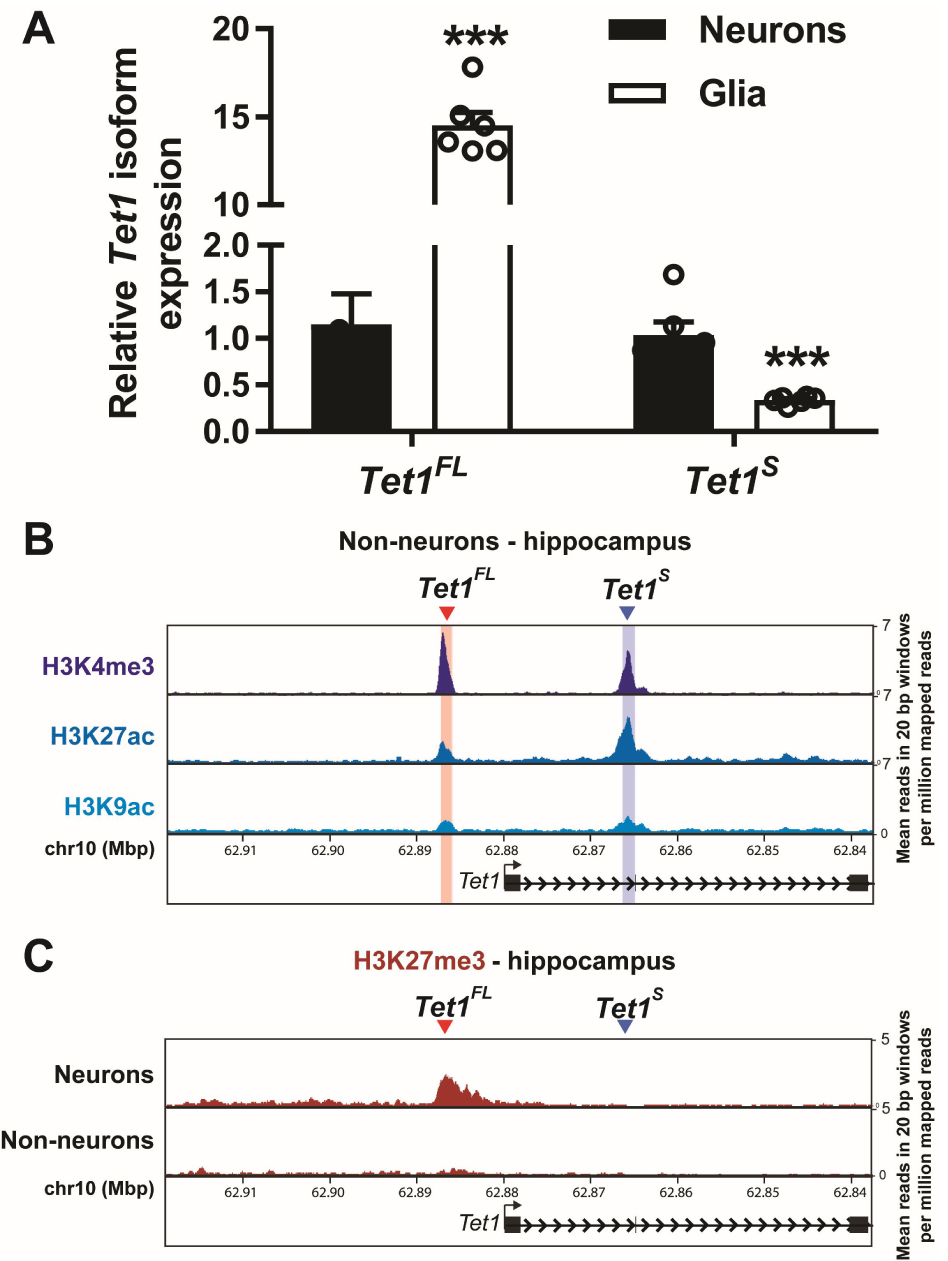
*Tet1* isoform transcript usage differs between neurons and non-neuronal cells in the brain. ***A***, qRT-PCR analysis of *Tet1*^*FL*^ and *Tet1*^*S*^ isoform expression levels in primary hippocampal glial cultures relative to hippocampal neuron cultures. *** p < 0.001 (unpaired two-tailed *t* test). n = 6. Data represent mean ± SEM. ***B***, Mean normalized H3K4me3, H3K27ac, and H3K9ac ChIP-seq signals in hippocampal non-neuronal cells (NeuN-) at the *Tet1* gene locus. ***C***, Comparison of mean normalized H3K27me3 ChIP-seq signals in hippocampal neurons (NeuN+) and non-neuronal (NeuN-) at the *Tet1* gene locus. Data for both ***B*** and ***C*** generated from Halder *et al*., 2015.

To explore these differences further, we compared the chromatin status of each *Tet1* isoform promoter in non-neuronal cells (NeuN-) to that of neurons using previously published ChIP-seq datasets (Halder et al., 2015). Similar to neurons, we found in NeuN-cells that both *Tet1* isoform promoters were co-enriched for H3K4me3, H3K9ac, and H3K27ac. However, in NeuN-cells, the largest H3K4me3 peak was located at the *Tet1*^*FL*^ promoter (Fig. 2*B*), whereas in neurons we observed the strongest enrichment at the *Tet1*^*S*^ promoter. In addition, we found that in neurons, the *Tet1*^*FL*^ promoter was marked by the repressive histone modification H3K27me3 (Fig. 2*C*). The presence of active (H3K4me3, H3K27ac, H3K9ac) and repressive (H3K27me3) histone marks at *Tet1*^*FL*^ exon 1 in neurons suggests that the *Tet1*^*FL*^ promoter is bivalent in these cells, which has been shown to keep genes expressed at low basal levels, poised for reactivation (Bernstein et al., 2006). However, it is important to note that while the ChIP-seq libraries we used are neuron-specific, they represent a heterogeneous collection of neuronal subtypes. Thus, the presence of bivalent histone marks at the *Tet1*^*FL*^ promoter may represent a characteristic of all neurons, or a mixture where some express and some silence *Tet1*^*FL*^. Regardless, our findings demonstrate that the *Tet1* isoform transcripts are differentially expressed in neurons and glia.

### *Tet1*^*S*^ transcript levels are downregulated in response to neuronal activity

We and others previously reported that total *Tet1* mRNA levels are decreased in response to neuronal activity (Kaas et al., 2013; Widagdo et al., 2014). To examine the contributions of the *Tet1*^*FL*^ and *Tet1*^*S*^ transcripts to these changes, hippocampal cultures were incubated for 1 or 4 h with KCl, bicuculline, or N-methyl-D-aspartate and glycine (NMDA/gly), and their expression levels were evaluated using qRT-PCR. We found that *Tet1*^*FL*^ mRNA levels were unaffected by any of the treatments (fold change: KCl-*FL*, F (2, 45) = 0.6, p = 0.5513, n = 16; bic-*FL*, F (2, 15) = 0.25, p = 0.7788, n = 6; NMDA/Gly-*FL*, (F (2, 33) = 0.091, n = 12; One Way ANOVA) (Fig. 3*A*). In contrast, *Tet1*^*S*^ mRNA levels were significantly decreased at both the 1 h and 4 h time points after KCl stimulation (fold change: F (2, 45) = 15, p < 0.0001, One Way ANOVA; veh, 1 ± 0.034 vs. 1 h, 0.76 ± 0.048, p < 0.0001, n = 16; veh, 1 ± 0.034 vs. 4h, 0.75 ± 0.024, p < 0.0001, n = 16; Dunnett’s *post hoc*) and at the 4 h time point after treatment with either bicuculline (fold change: F (2, 15) = 15, p = 0.0003, One Way ANOVA, veh, 1 ± 0.048 vs. 4h, 0.64 ± 0.041, p = 0.0002, n = 6, Dunnett’s *post hoc*) or NMDA/gly (fold change: F (2, 33) = 21, p < 0.0001, One Way ANOVA; veh, 1 ± 0.025 vs. 4h, 0.58 ± 0.045, p < 0.0001, n = 12, Dunnett’s *post hoc*), suggesting that its expression is regulated by activity- and NMDA receptor-dependent mechanisms (Fig. 3*B*). We confirmed that each of these treatments significantly increased the expression levels of the immediate early gene (IEG) *Activity regulated cytoskeleton associated protein* (*Arc*), as expected (fold-change: KCl-*Arc* - F (2, 44) = 6.034, p = 0.0048, One Way ANOVA; Veh, 1 ± 0.063 vs. 1h, 3.3 ± 0.4, p = 0.0388, Veh, 1 ± 0.063 vs. 4h, 4.2 ± 1.1, p = 0.0030, Dunnett’s *post hoc*; bic-*Arc* - F (2, 15) = 113.4, p < 0.0001, One Way ANOVA, Veh, 1± 0.097 vs. 1h, 7.1 ± 0.4, p < 0.0001, Dunnett’s *post hoc*; NMDA/Gly-*Arc* - F (2, 33) = 27.02, p < 0.0001, One Way ANOVA, Veh, 1.1 ± 0.064 vs. 1h, 4.9 ± 0.68, p < 0.0001, Dunnett’s *post hoc*) (Fig. 3*C*).

**Figure 3.**
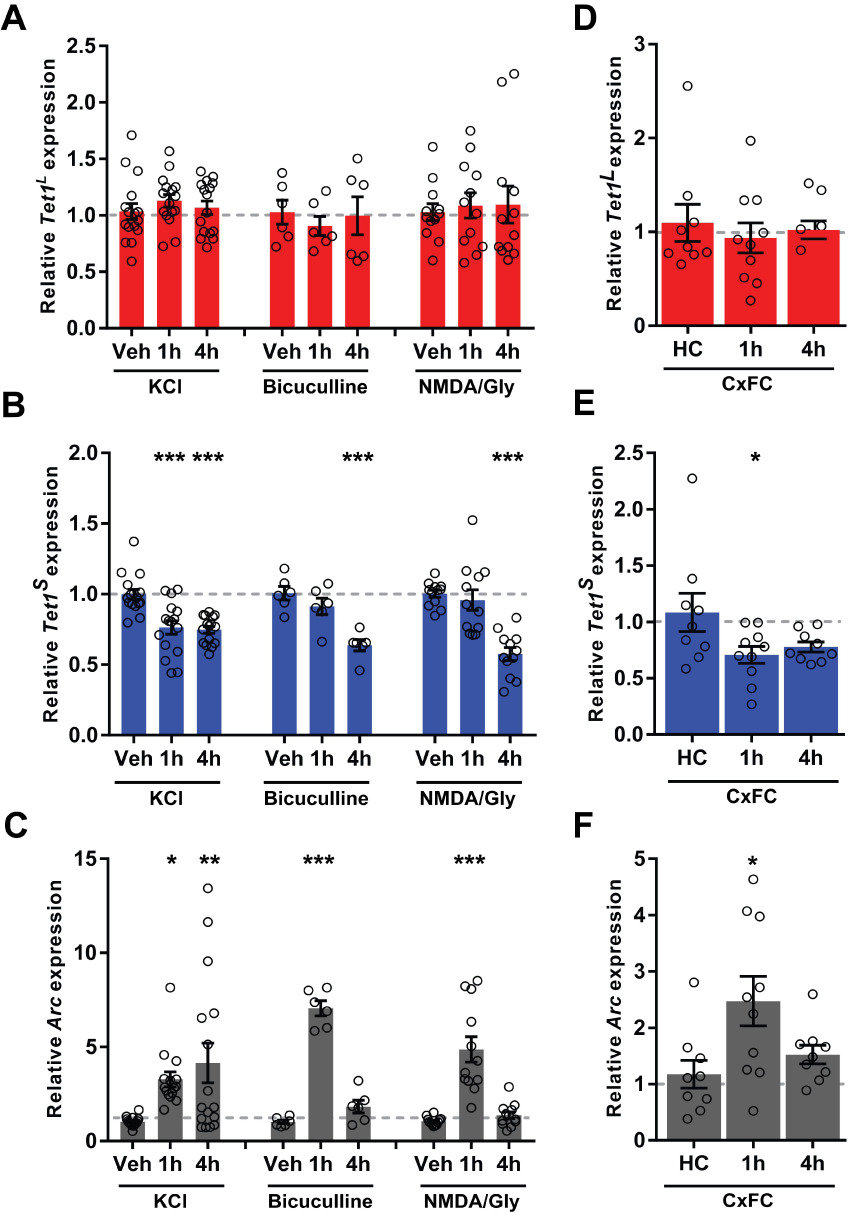
Neuronal activity-dependent downregulation of *Tet1*^*S*^ transcript levels. qRT-PCR analysis of ***A***, *Tet1*^*FL*^ ***B***, *Tet1*^*S*^ and ***C***, *Arc* expression levels in primary hippocampal neurons 1 and 4 h after stimulation with 25 mM KCl, 50 µM Bicuculline or a combination of 10 µM NMDA + 2 µM glycine. * p < 0.05, ** p < 0.01, *** p < 0.001 (Dunnett’s *post hoc* vs. vehicle). p < 0.001 (One Way ANOVA). KCl, n = 16, Bicuculline, n = 6, NMDA + Glycine, n = 12. Data represent mean ± SEM. qRT-PCR analysis of ***D***, *Tet1*^*FL*^ ***E***, *Tet1*^*S*^ *and* ***F***, *Arc* expression levels in hippocampal area CA1 1 and 4 h after CxFC (3 context-shock pairings, 0.75mA, 2s). * p < 0.05 (Dunnett’s *post hoc* vs. home cage (HC)). p < 0.05 (One Way ANOVA). HC, n = 9 mice, 1 h, n = 10 mice, 4 h, n = 9 mice. Data represent mean ± SEM.

Next, we measured *Tet1*^*FL*^ and *Tet1*^*S*^ expression levels in hippocampal area CA1 tissue after contextual fear conditioning (CxFC) to examine whether both isoforms responded similarly to neuronal activity induced *in vivo* during memory formation. Again, we observed that *Tet1*^*FL*^ expression levels remained unchanged (fold change: F (2, 25) = 0.2666, p = 0.7681, n = 9-10, One Way ANOVA) (Fig. 3*D*), whereas *Tet1*^*S*^ levels were significantly down-regulated 1 h after training (fold change: F (2, 25) = 3.399, p = 0.0494, One Way ANOVA, hc, 1.1 ± 0.17, vs. 1h, 0.71 ± 0.075, p = 0.0374, n = 9-10, Dunnett’s *post hoc*) (Fig. 3*E*). As in the primary cultures, *Arc* expression was significantly induced in the hippocampus during our CxFC paradigm (CxFC-*Arc -* F (2, 25) = 4.554, p = 0.0206, One Way ANOVA, hc, 1.2 ± 0.25 vs. 1h, 2.5 ± 0.44, p = 0.0146, Dunnett’s *post hoc*) (Fig. 3*F*). Together, these results show that transcript levels of the *Tet1*^*S*^ gene are generally down-regulated in response to hippocampal neuron stimulation, suggesting that under basal conditions the isoform may act as a molecular restraint on activity-dependent processes in neurons.

### Individual manipulation of *Tet1*^*FL*^ and *Tet1*^*S*^ expression in neurons

The differential regulation of *Tet1* isoform expression following neuronal activation suggests that that *Tet1*^*FL*^ and *Tet1*^*S*^ may have unique cell-specific functions.

However, prior studies globally manipulated total *Tet1* expression levels, providing limited insight into the cell-specific functions of the *Tet1* isoforms. Therefore, we developed genetic tools to selectively manipulate *Tet1*^*FL*^ and *Tet1*^*S*^ expression levels in hippocampal neurons, both in culture and *in vivo*. To accomplish this, we designed sequence-programmable Transcription Activator Like Effectors (TALEs) to selectively target *Tet1*^*FL*^ or *Tet1*^*S*^ due to their previously reported high target specificity, cell type-specific expression, and small size compatible with *in vivo* delivery using a signal AAV virus (Konermann et al., 2013; Mendenhall et al., 2013; Juillerat et al., 2014; Polstein et al., 2015). HA-tagged TALEs were designed to specifically bind to DNA sequences at each *Tet1* isoform promoter and either repress transcription (TALE-SID4X, 4 copies of the Sin3a Interacting) or serve as a target sequence-specific control (TALE-NFD, no functional domain) (Konermann et al., 2013; Choi et al., 2014) (Fig. 4*A, B*). TALE expression in these modified AAV vectors was placed under the control of the human synapsin promoter, which drives expression only in neurons (Kügler et al., 2003). We found that expression of either control TALE in Neuro2a (N2a) cells, hereafter referred to as *Tet1*^*FL*^-NFD and *Tet1*^*S*^-NFD, did not alter the expression of the *Tet1* isoforms relative to mock-transfected cells, suggesting that TALE binding to these promoter regions without an effector domain did not sterically hinder transcription (fold change: mock, 1 ± 0.095 vs. *Tet1*^*FL*^-NFD, 1 ± 0.061, t_(6)_ = 0.069, p = 0.9473; mock, 1 ± 0.053 vs. *Tet1*^*S*^-NFD, 1.1 ± 0.067, t_(6)_ = 0.68, p = 0.5242, n = 4 all groups, unpaired two-tailed *t* test) (Fig. 4*C, D*). Conversely, expression of the TALE repressors, here after referred to as *Tet1*^*FL*^-SID4X and *Tet1*^*S*^-SID4X, significantly inhibited the expression of *Tet1*^*FL*^ and *Tet1*^*S*^ in N2a cells, respectively. Importantly, neither TALE repressor affected the expression of the opposite isoform, indicating isoform-specific targeting. (fold change: *FL-*F (2, 9) = 26, p = 0.0002, One Way ANOVA, *Tet1*^*FL*^-NFD, 1.1 ± 0.044 vs. *Tet1*^*FL*^-SID4X, 0.42 ± 0.021, p = 0.0006, *Tet1*^*FL*^-NFD, 1.1 ± 0.044 vs. *Tet1*^*S*^-SID4X, 1.2 ± 0.14, p = 0.5668, n = 4 all groups, Dunnett’s *post hoc*; *S-*F (2, 9) = 17, p = 0.0008, One Way ANOVA, *Tet1*^*S*^-NFD, 1 ± 0.061 vs. *Tet1*^*S*^-SID4X, 0.15 ± 0.0039, p = 0.0009, *Tet1*^*S*^-NFD, 1 ± 0.061 vs. *Tet1*^*FL*^-SID4X, 0.91 ± 0.18, p = 0.7794, n = 4 all groups, Dunnett’s *post hoc*) (Fig. 4*E, F*). To test the function of the *Tet1* isoform-specific TALEs in primary cells, we packaged each construct into AAV1 viral particles and and evaluated their efficacy and specific in transduced hippocampal neurons. Owing to their cell type specificity, immunocytochemistry revealed that the TALE constructs only expressed in cells positive for the neuronal marker NeuN (Fig. 4*G*). Similar to their effects in N2a cells, transduction of primary hippocampal neurons with AAV1-*Tet1*^*FL*^-SID4X and *Tet1*^*S*^-SID4X led to a significant reduction in the expression levels of the intended *Tet1* isoform target, without affecting the opposite transcript (fold change: *FL-*F (2, 24) = 7.549, p = 0.0029, One Way ANOVA, *Tet1*^*FL*^-NFD, 1 ± 0.11 vs. *Tet1*^*FL*^-SID4X, 0.45 ± 0.073, p = 0.0378, *Tet1*^*FL*^-NFD, 1 ± 0.11 vs. *Tet1*^*S*^-SID4X, 1.4 ± 0.26, p = 0.3153, n = 9 all groups, Dunnett’s *post hoc*; *S-*F (2, 24) = 35.42, p < 0.0001, One Way ANOVA, *Tet1*^*S*^-NFD, 1 ± 0.02 vs. *Tet1*^*S*^-SID4X, 0.17 ± 0.033, p < 0.0001, *Tet1*^*S*^-NFD, 1 ± 0.02 vs. *Tet1*^*FL*^-SID4X, 0.92 ± 0.13, p = 0.6628, n = 9 all groups, Dunnett’s *post hoc*) (Fig. 4*H, I*). Finally, we measured transcript levels of the IEGs *Neuronal PAS domain protein 4* (*Npas4*), *Arc*, and *Early growth response 1* (*Egr1*) in primary hippocampal neurons transduced with AAV1-*Tet1*^*FL*^-SID4X or -*Tet1*^*S*^-SID4X because *Tet1* was previously shown to regulate their expression in the brain (Kaas et al., 2013; Rudenko et al., 2013; Kumar et al., 2015; Towers et al., 2018). We found that SID4X-mediated repression of either *Tet1* isoform led to a significant reduction in the expression of all three genes compared to NFD controls (fold change: *Tet1*^*FL*^-NFD vs. *Tet1*^*FL*^-SID4X-*Npas4*, 1.1 ± 0.19 vs. 0.38 ± 0.041, t_(16)_ = 3.7, n = 9, p = 0.002, *Arc*, 1.1 ± 0.14, vs.0.35 ± 0.04, t_(16)_ = 5, n = 9, p < 0.0001, *Egr1*, 1 ± 0.027, vs. 0.5 ± 0.043, t^(12)^ = 10, n =7, p < 0.0001; *Tet1*^*S*^-NFD vs. *Tet1*^*S*^-SID4X-*Npas4*, 1.1 ± 0.22 vs. 0.32 ± 0.057 t_(16)_ = 3.6, n = 9, p = 0.0023, *Arc*, 1.2 ± 0.23 vs. 0.51 ± 0.12, t_(16)_ = 2.6, n = 9, p = 0.021, *Egr1*, 1 ± 0.032 vs. 0.35 ± 0.014, t_(12)_ = 19, n = 7, p < 0.0001, unpaired two tailed *t* test) (Fig. 4*J, K*). Together, these results demonstrate that our modified TALE tools significantly repress the transcription of each individual *Tet1* isoform, are neuron-specific, and result in changes in the expression of genes previously shown to be targets of TET1 in the central nervous system.

**Figure 4.**
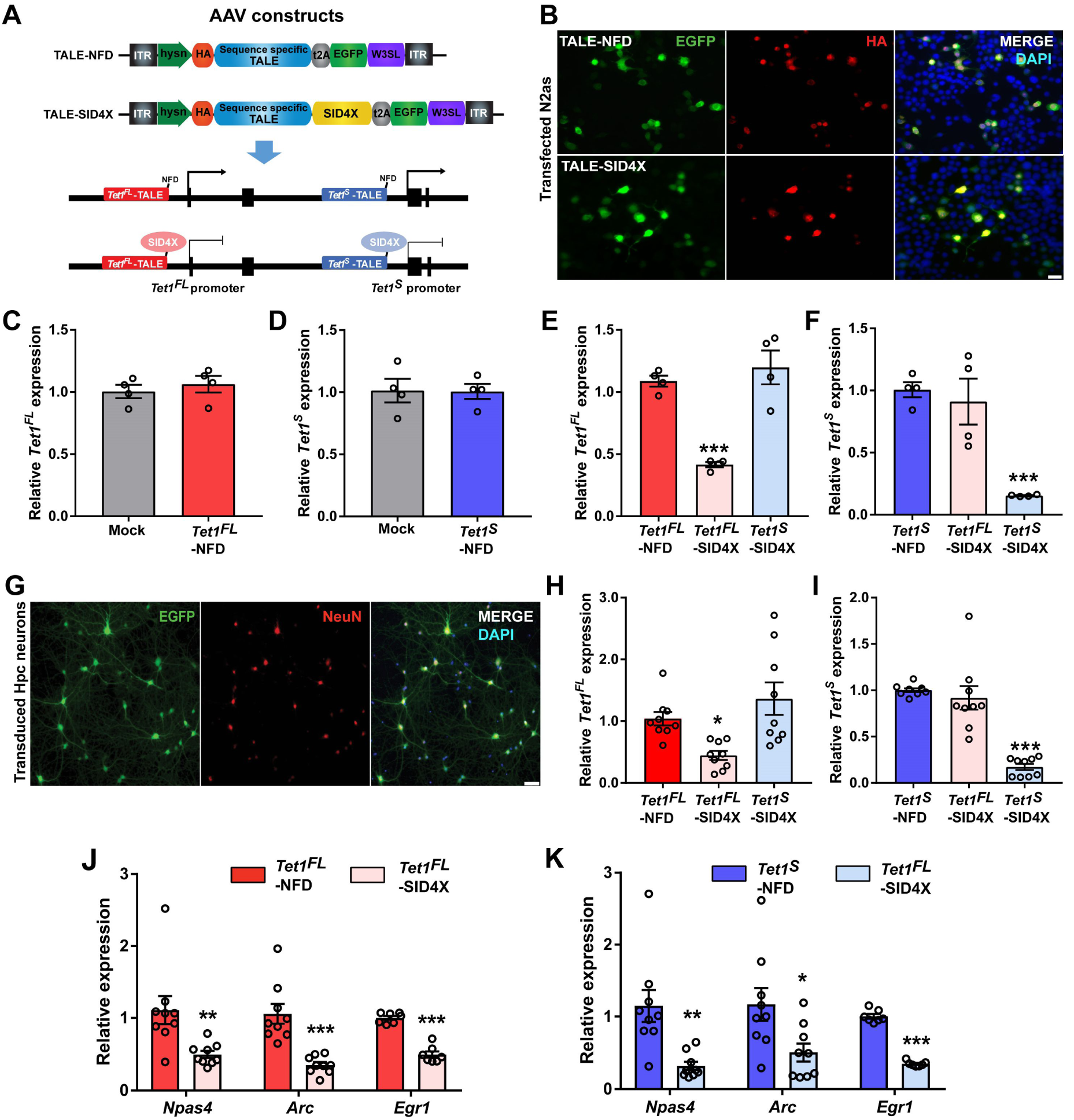
Construction and validation of synthetic transcription factors to repress endogenous *Tet1* isoform expression in neurons. ***A***, Top, Illustration of neuron-specific TALE-construct modifications to increase target sequence capacity. ITR, inverted terminal repeats; h*SYN*, human Synapsin promoter; HA, Human influenza hemagglutinin tag; TALE, transcriptional activator like effector; t2A, thosea asigna virus 2A (self-cleaving peptide); SID4X, 4 copies of the Sin3a Interacting Domain; EGFP, enhanced green fluorescent protein; W3SL, truncated woodchuck hepatitis posttranscriptional regulatory element and polyadenylation signal cassette (Choi et al., 2014). Bottom, strategy to transcriptionally repress *Tet1*^*FL*^ and *Tet1*^*S*^ using TALE-SID4X constructs. ***B***, Representative images of GFP and HA immunostaining in N2a cells expressing TALE-NFD, or -SID4X constructs. Scale bar, 20 µM. ***C***, qRT-PCR analysis of *Tet1*^*FL*^ expression levels in mock and *Tet1*^*FL*^-NFD transfected N2a cells. n = 4. ***D***, qRT-PCR analysis of *Tet1*^*S*^ expression levels in mock and *Tet1*^*S*^*-*NFD transfected N2a cells. n = 4. ***E***, qRT-PCR analysis of *Tet1*^*FL*^ expression levels in N2a cells transfected with *Tet1*^*FL*^-NFD, *Tet1*^*FL*^-SID4X or *Tet1*^*S*^-SID4X. n = 4. ***F***, qRT-PCR analysis of *Tet1*^*S*^ expression levels in N2a cells transfected with *Tet1*^*S*^-NFD, *Tet1*^*FL*^-SID4X or *Tet1*^*S*^-SID4X. n = 4. ***G***, Representative EGFP and NeuN immunostaining of primary hippocampal cultures 5 days after transduction with AAV1-TALE constructs. Scale bar, 50 µM. ***H***, qRT-PCR analysis of *Tet1*^*FL*^ expression levels in primary hippocampal neurons transduced with AAV1-*Tet1*^*FL*^-NFD, -*Tet1*^*FL*^-SID4X or -*Tet1*^*S*^-SID4X. n = 9. ***I***, qRT-PCR analysis of *Tet1*^*S*^ expression levels in primary hippocampal neurons transduced with AAV1-*Tet1*^*S*^-NFD, -*Tet1*^*FL*^-SID4X or -*Tet1*^*S*^-SID4X. n = 9. ***J***, qRT-PCR analysis of *Npas4, Arc* and *Egr1* expression levels in primary hippocampal neurons transduced with AAV1-*Tet1*^*FL*^-NFD and -*Tet1*^*FL*^-SID4X. n = 9. ***K***, qRT-PCR analysis of *Npas4, Arc* and *Egr1* expression levels in primary hippocampal neurons transduced with *Tet1*^*S*^-NFD and *Tet1*^*S*^-SID4X. n = 9. * p < 0.05, ** p < 0.01, *** p < 0.001 (Dunnett’s *post hoc*). p < 0.05 (One Way ANOVA). All data represent mean ± SEM.

### *Tet1*^*FL*^ and *Tet1*^*S*^ regulate unique subsets of the neuronal transcriptome

To investigate the effects of each individual *Tet1* isoform on neuronal gene expression, we infected primary hippocampal neurons with AAV1-TALEs and performed an unbiased, transcriptome-wide RNA-seq analysis on control and *Tet1* isoform-depleted samples. We found that despite its low transcript levels, acute repression of *Tet1*^*FL*^ caused widespread transcriptional changes in neurons. Using a cutoff greater than -/+ 0.2 log2FC and an FDR < 0.05, we identified more than 6,000 differentially expressed genes (DEGs; Fig. 5*A*, top, Extended data table 5-1). Gene Ontology (GO) analysis revealed that *Tet1*^*FL*^*-*modulated mRNAs functioned in a wide assortment of biological processes (BP) and Kyoto Encyclopedia of Genes and Genomes pathways (KEGG, Fig. 5*A*, bottom, Extended data table 5-2, 5-3). In both sets of analyses, genes downregulated in response to *Tet1*^*FL*^ repression were generally enriched for neuron-associated functions, such as ion transport, learning, long-term synaptic potentiation and the synaptic vesicle cycle (BP terms and KEGG terms: ion transport-3.3 × 10^−20^, learning-3 × 10^−12^, long-term synaptic potentiation-7.8 × 10^−8^, synaptic vesicle cycle-3.2 × 10^−7^; Benjamini–Hochberg adjusted p values) (Fig. 5*A*, bottom left). In contrast, upregulated genes were enriched for BP terms associated with the cell cycle, DNA damage, and immune function (6.3 × 10^−25^, 1.6 × 10^−14^, 1.3 × 10^−9^, respectively; Benjamini–Hochberg adjusted p values) as well as several KEGG pathway categories related to cancer pathways, extracellular matrix (ECM) receptor interactions, and NF-κB signaling (3.5 × 10^−15^, 1.1 × 10^−11^, 7.3 × 10^−10^, respectively; Benjamini–Hochberg adjusted p values)(Fig. 5*A*, bottom right). In the case of *Tet1*^*S*^, we found that its repression led to less than a quarter of the number of DEGs compared to *Tet1*^*FL*^ (Fig. 5*B*, top, Extended data table 5-4). Genes upregulated following *Tet1*^*S*^ repression were enriched for some of the same BP categories as *Tet1*^*FL*^, but also included terms associated with ribosomal biogenesis, methylation, and covalent chromatin modifications (5.1 × 10^−4^, 2 × 10^−3^, 3.7 × 10^−3^, 3.2 × 10^−7^, respectively; Benjamini–Hochberg adjusted p values) (Fig. 5*B*, bottom right, Extended data table 5-5). Notably, downregulated genes associated with *Tet1*^*S*^ were not significantly enriched in any pathway or GO category (Extended data table 5-6).

**Figure 5.**
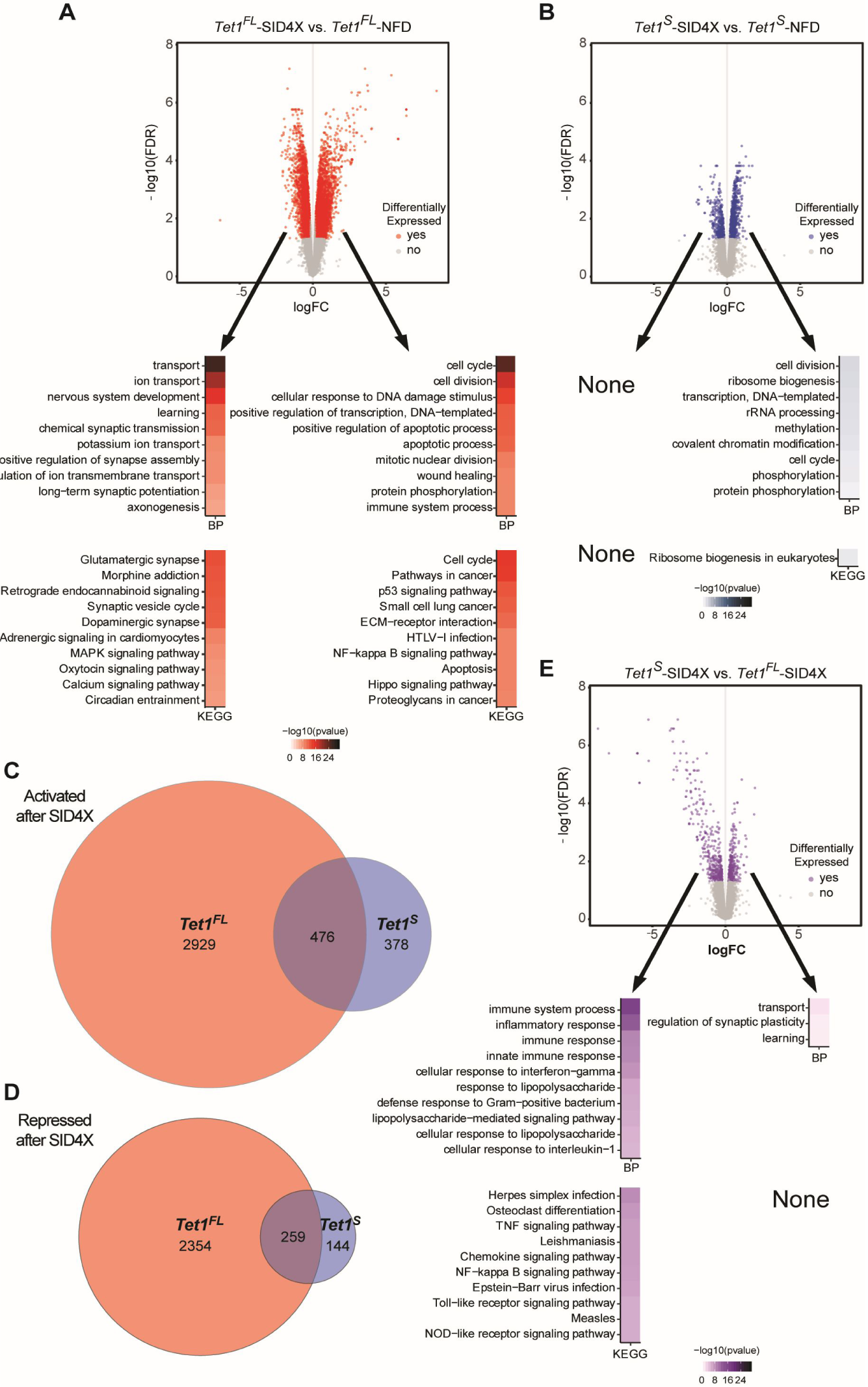
Transcriptomic analysis of hippocampal neuron cultures depleted of *Tet1*^*FL*^ and *Tet1*^*S*^. ***A***, Top: Volcano plot analysis of genome wide RNA-seq data comparing AAV1-*Tet1*^*FL*^-NFD and *Tet1*^*FL*^-SID4X transduced hippocampal neurons (+/- 0.2 log2FC, FDR < 0.05). n = 3 biological replicates. Bottom: 10 most statistically significant gene ontology biological processes (BP) and Kyoto Encyclopedia of Genes and Genomes (KEGG) enrichment terms associated with up- and downregulated DEGs in *Tet1*^*FL*^-SID4X versus *Tet1*^*FL*^-NFD transduced hippocampal neurons. p < 0.05 (Benjamini–Hochberg corrections). ***B***, Volcano plot analysis of genome wide RNA-seq data comparing AAV1-*Tet1*^*S*^-NFD and -*Tet1*^*S*^-SID4X transduced hippocampal neurons (+/- 0.2 log2FC, FDR < 0.05). n = 3 biological replicates. Bottom: Statistically significant gene ontology BP and KEGG enrichment terms of upregulated DEGs in AAV1-*Tet1*^*S*^-SID4X versus -*Tet1*^*S*^-NFD transduced hippocampal neurons. None; no statistically significant GO terms identified. p < 0.05 (Benjamini–Hochberg corrections). ***C***, Summary of the numbers of overlapping and non-overlapping significantly upregulated genes in AAV1-*Tet1*^*S*^-SID4X versus -*Tet1*^*FL*^-SID4X transduced hippocampal neurons relative to their respective NFD controls. ***D***, Summary of significantly downregulated genes. ***E***, Top: Volcano plot analysis of genome wide RNA-seq data comparing AAV1-*Tet1*^*S*^-SID4X and -*Tet1*^*FL*^-SID4X transduced hippocampal neurons (+/- 0.2 log2FC, FDR < 0.05). n = 3 biological replicates. Bottom: 10 most statistically significant gene ontology BP and KEGG enrichment terms associated with up- and downregulated DEGs in AAV1-*Tet1*^*S*^-SID4X versus -*Tet1*^*FL*^-SID4X transduced hippocampal neurons. p < 0.05 (Benjamini–Hochberg corrections).

To further explore differences between *Tet1*^*FL*^ and *Tet1*^*S*^-mediated transcriptional regulation, we performed a direct comparison of *Tet1*^*S*^*-*SID4X to *Tet1*^*FL*^*-*SID4X associated DEGs. Slightly over half of activated genes (Fig. 5*C*, Extended data table 5-7) and repressed genes (Fig. 5*D*, Extended data table 5-7) after modulation of the short isoform were also changed after suppression of the long isoform. Through this comparison, however, we found that significant differentially expressed genes modulated by changes in the expression of each isoform did not entirely overlap, suggesting these two isoforms do not functionally identical. This prompted us to do a direct comparison of the SID4X conditions, and this further statistical analysis (Extended data table 5-8) revealed that DEGs expressed at lower levels in the *Tet1*^*S*^-SID4X dataset, relative to *Tet1*^*FL*^-SID4X, were generally involved in immune system regulation (BP terms; Immune system process - 1.7 × 10^−17^, inflammatory response - 2.2 × 10^−13^, innate immune response - 1.6 × 10^−9^: KEGG terms; TNF signaling pathway - 4.9 × 10^−8^, Nf-κB signaling pathway - 6.5 × 10^−^_8_; Benjamini– Hochberg adjusted p values) (Fig. 5*E*, bottom left, Extended data table 5-9). Genes more abundantly expressed when *Tet1*^*S*^ was repressed relative to *Tet1*^*FL*^, functioned in transport, regulation of synaptic plasticity, and learning (1.2 × 10^−2^, 3.3 × 10^−2^, 3.7 × 10^−2^, respectively; Benjamini–Hochberg adjusted p values) (Fig. 5*E*, bottom right, Extended data table 5-10). We interpret these data to mean that while overlaps exist, the acute repression of *Tet1*^*FL*^ aberrantly activates inflammatory response pathways, while *Tet1*^*S*^ does not. Moreover, relative to *Tet1*^*FL*^, the acute suppression of *Tet1*^*S*^ elicits higher expression of genes involved in synaptic plasticity.

### Acute repression of *Tet1*^*FL*^ and *Tet1*^*S*^ expression has opposing effects on synaptic transmission

To compare the effects of selectively disrupting *Tet1*^*FL*^ or *Tet1*^*S*^ expression on excitatory synaptic transmission, we infected hippocampal neurons with AAV1-TALEs and measured the amplitude and frequency of miniature excitatory postsynaptic currents (mEPSCs) in EGFP positive control and *Tet1* isoform-depleted cells. SID4X-mediated repression of *Tet1*^*FL*^ significantly increased both mEPSC amplitude and frequency compared to controls (Fig. 6*A, B, C*), as reflected by significant rightward shifts in the cumulative probability distributions of both measurements (amplitude; *D =* 0.1044, p = 0.012: frequency; *D =* 0.2325, p < 0.0001, *Tet1*^*FL*^-NFD, n = 19, *Tet1*^*FL*^-SID4X, n = 20; Kolmogorov-Smirnov test). In contrast, *Tet1*^*S*^ repression had no effect on mEPSC amplitude (Fig. 6*D, E*), but significantly reduced mEPSC frequency (Fig. 6*D, F*), illustrated by a significant leftward shift in the cumulative probability distribution (amplitude; *D* = 0.0659, p = 0.151: frequency; *D =* 0.3230, p < 0.0001, *Tet1*^*S*^- NFD, n = 20, *Tet1*^*S*^-SID4X, n = 23; Kolmogorov-Smirnov test). Overall, these data suggest that the acute and selective repression of *Tet1* isoform expression differentially regulates excitatory synaptic transmission.

**Figure 6.**
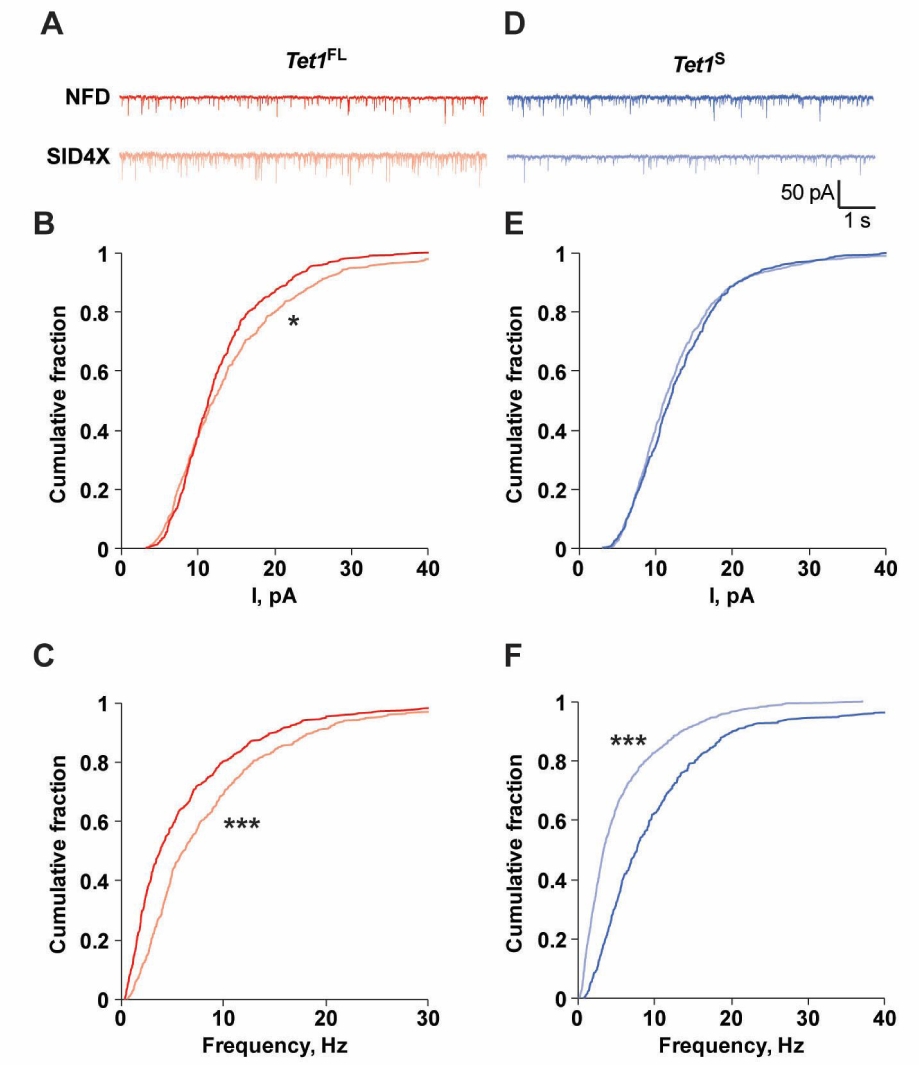
*Tet1*^*FL*^ and *Tet1*^*S*^ repression alters excitatory synaptic transmission. ***A***, Representative mEPSC traces from hippocampal neurons transduced with AAV1-*Tet1*^*FL*^-NFD and -*Tet1*^*FL*^-SID4X, Calibration: 1 s, 50 pA. ***B***, Cumulative probability plot of mEPSC amplitude in hippocampal neurons transduced with AAV1-*Tet1*^*FL*^-NFD and -*Tet1*^*FL*^-SID4X. *p < 0.05 (Kolmogorov-Smirnov test). n = 19-20. ***C***, Cumulative probability plot of mEPSC frequency in hippocampal neurons transduced with AAV1-*Tet1*^*FL*^-NFD and -*Tet1*^*FL*^-SID4X. ***p < 0.001 (Kolmogorov-Smirnov test). n = 19-20. ***D***, Representative mEPSC traces from hippocampal neurons transduced with AAV1-*Tet1*^*S*^-NFD and -*Tet1*^*S*^-SID4X, Calibration: 1 s, 50 pA. ***E***, Cumulative probability plots of mEPSC amplitude in hippocampal neurons transduced with AAV1-*Tet1*^*S*^-NFD and -*Tet1*^*S*^-SID4X. p > 0.05 (Kolmogorov-Smirnov test). n = 20-23. ***F***, Cumulative probability plot of mEPSC frequency in hippocampal neurons transduced with AAV1-*Tet1*^*FL*^-NFD and -*Tet1*^*FL*^-SID4X. ***p < 0.001 (Kolmogorov-Smirnov test). n = 20-23.

### *Tet1*^*FL*^ and *Tet1*^*S*^ differentially regulate hippocampal-dependent memory

We next assessed the cognitive effects of selectively inhibiting *Tet1*^*FL*^ and *Tet1*^*S*^ expression in the dorsal hippocampus (dHPC) using our neuron- and isoform-specific molecular tools. We found that stereotaxic injection of *Tet1* isoform-specific AAV1-TALEs into the dorsal hippocampus led to widespread EGFP expression throughout the CA1-CA3 subfields after two weeks (Fig. *7A*). In addition, the expression levels of *Tet1*^*FL*^ and *Tet1*^*S*^ were significantly reduced by AAV1-*Tet1*^*FL*^-SID4X and -*Tet1*^*S*^-SID4X transduction relative to AAV1-*Tet1*^*FL*^-NFD and *Tet1*^*S*^-NFD controls, indicating that the molecular tools were effective *in vivo* (fold change: *Tet1*^*FL*^-NFD, 1.1 ± 0.14 vs. *Tet1*^*FL*^-SID4X, 0.73 ± 0.051, t _(14)_ = 2.6, p = 0.02, n = 8: *Tet1*^*S*^-NFD, 1.1 ± 0.063 vs. *Tet1*^*S*^-SID4X, 0.57 ± 0.044, t _(14)_ = 6.2, p < 0.0001, n = 8, unpaired two-tailed *t* test (Fig. 7*B, C*).

**Figure 7.**
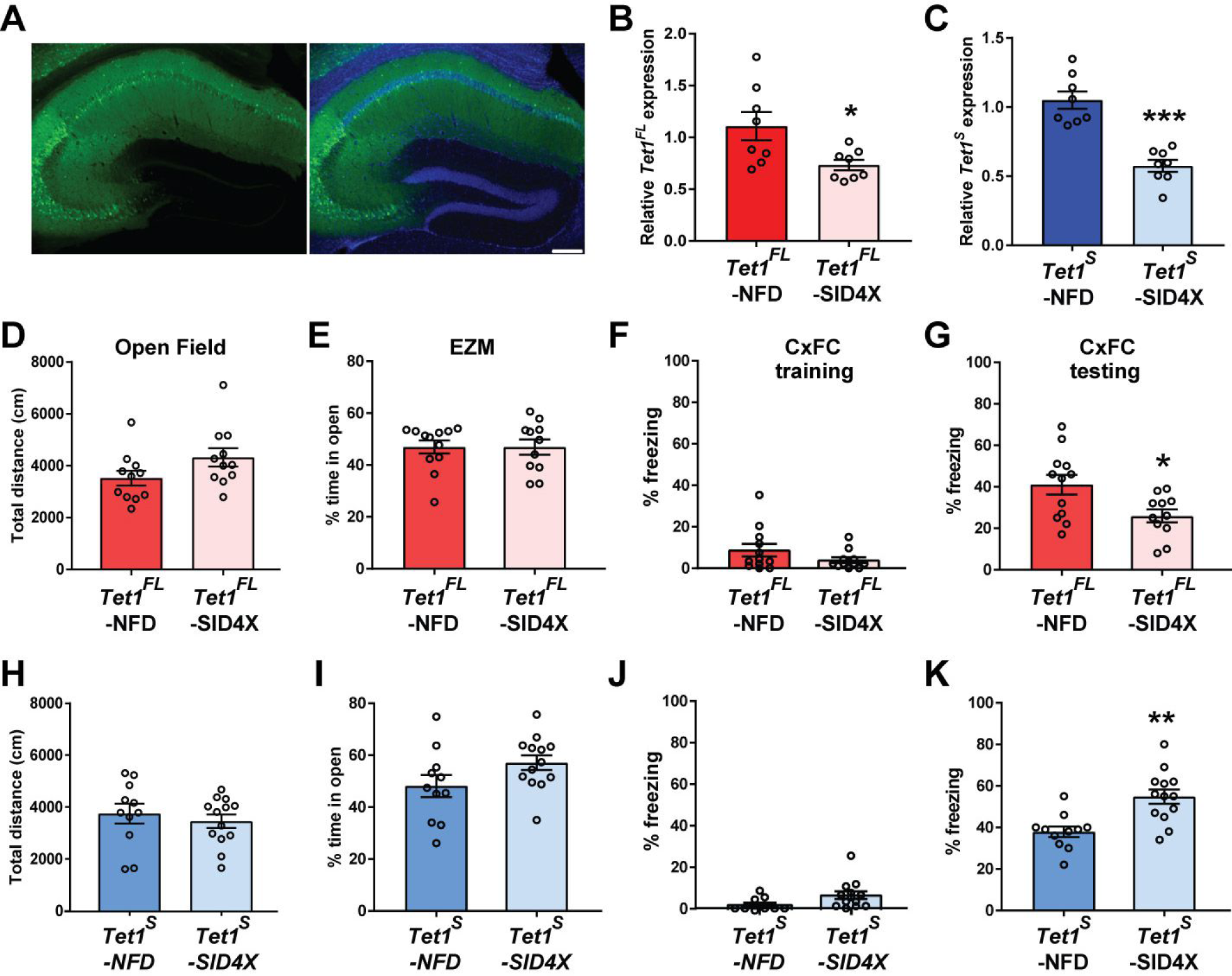
Neuron-specific *Tet1*^*FL*^ and *Tet1*^*S*^ repression differentially affect hippocampal-dependent memory formation. ***A***, Left, Representative EGFP immunostaining in the dorsal hippocampus 2 weeks after AAV1-mediated transduction of *Tet1* isoform-specific TALE constructs. Right, Merge of EGFP immunostaining and DAPI. Scale bar, 100 µm. ***B***, qRT-PCR analysis of *Tet1*^*FL*^ expression levels in the dorsal hippocampus two weeks after AAV1-*Tet1*^*FL*^-NFD and *Tet1*^*FL*^-SID4X transduction. * p < 0.05 (unpaired two-tailed *t* test). n = 8. ***C***, qRT-PCR analysis of *Tet1*^*S*^ expression levels in the dorsal hippocampus two weeks after AAV1-*Tet1*^*S*^-NFD and -*Tet1*^*S*^-SID4X transduction. *** p < 0.001 (unpaired two-tailed *t* test). n = 8. ***D***, Total distance traveled (cm) during a 30 min Open field exploration test in AAV1-*Tet1*^*FL*^-NFD and *Tet1*^*FL*^-SID4X transduced mice. n = 11-12. ***E***, Total distance traveled (cm) during a 30 min Open field exploration test in AAV1-*Tet1*^*FL*^-NFD and *Tet1*^*FL*^-SID4X transduced mice. n = 11-13. ***F***, Percent time spent freezing in AAV1-*Tet1*^*FL*^-NFD and *Tet1*^*FL*^-SID4X transduced mice during a 3.5 min CxFC session. n = 11-12. Percent time in the open arms during 5 min of exploration in the EZM in AAV1-*Tet1*^*FL*^-NFD and *Tet1*^*FL*^-SID4X transduced mice. n = 11-12. ***G***, Percent time spent freezing in AAV1-*Tet1*^*FL*^-NFD and *Tet1*^*FL*^-SID4X transduced mice during a 5 min CxFC test 24 h after training. * p < 0.05 (unpaired two-tailed *t* test). n = 11-12. ***H***, Total distance traveled (cm) during a 30 min Open field exploration test in AAV1-*Tet1*^*S*^-NFD and *Tet1*^*S*^-SID4X transduced mice. n = 11-12. ***I***, Percent time in the open arms during 5 min of exploration in the EZM in AAV1-*Tet1*^*S*^-NFD and *Tet1*^*S*^-SID4X transduced mice. n = 11-13. ***J***, Percent time spent freezing in AAV1-*Tet1*^*S*^-NFD and *Tet1*^*S*^-SID4X transduced mice during a 3.5 min CxFC session. n = 11-13. ***K***, Percent time spent freezing in AAV1-*Tet1*^*S*^-NFD and *Tet1*^*S*^-SID4X transduced mice during a 5 min CxFC test 24 h after training. ** p < 0.01 (unpaired two-tailed *t* test). n = 11-13. All data represent mean ± SEM.

We next tested whether transcriptional repression of *Tet1*^*FL*^ and *Tet1*^*S*^ in the dHPC led to any changes in locomotion, anxiety, or cognition. In both the open field and elevated zero maze (EZM) tests, there were no significant differences between mice infected with virus to silence either isoform compared to their controls (Open field, total distance (cm); *Tet1*^*FL*^-NFD, 3520 ± 283, n = 12 vs. *Tet1*^*FL*^-SID4X, 4322 ± 352, n = 11, t_(21)_ = 0.64, p = 0.5262; *Tet1*^*S*^-NFD, 3746 ± 379, n = 11 vs. *Tet1*^*S*^-SID4X, 3459 ± 255, n = 13, t_(20)_ = 1.8, p = 0.0908: EZM, percent time in open; *Tet1*^*FL*^-NFD, 47 ± 2.5%, n = 12 vs. *Tet1*^*FL*^-SID4X, 47 ± 3%, n = 11, t_(21)_ = 0.0041, p = 0.9968; *Tet1*^*S*^-NFD, 48 ± 4.2%, n = 11 vs. *Tet1*^*S*^-SID4X, 57 ± 2.8%, n = 13, t_(22)_ = 1.8, p = 0.838; unpaired two-tailed *t* test), suggesting that manipulation of *Tet1*^*FL*^ or *Tet1*^*S*^ in the dHPC does not affect general locomotion (Fig. 7*D, H)* or basal anxiety levels (Fig. 7*E, I*). Because previous studies examining the role of TET1 in memory formation were conducted in mice with disruptions that affect both isoforms (Rudenko et al., 2013; Zhang et al., 2013; Kumar et al., 2015; Towers et al., 2018), we next examined whether repression of either *Tet1*^*FL*^ or *Tet1*^*S*^ alone was sufficient to alter hippocampal-dependent memory formation. To test this, we trained mice using a moderate (see methods) CxFC paradigm and tested them 24 hours later, with the percentage of time spent freezing serving as an indirect measure of associative memory formation. We found no differences in the percentage of time freezing during training between *Tet1*^*FL*^-SID4X and *Tet1*^*FL*^*-*NFD mice (percent time freezing: *Tet1*^*FL*^*-*NFD, 8.1 ± 2.8%, n = 12 vs. *Tet1*^*FL*^-SID4X, 3.6 ± 1.3%, n = 11, t_(21)_ = 1.4, p = 0.1821; unpaired two-tailed *t* test), nor between the *Tet1*^*S*^-SID4X and *Tet1*^*S*^-NFD groups (percent time freezing: *Tet1*^*S*^*-*NFD, 3.6 ± 1.2%, n = 11 vs. *Tet1*^*S*^-SID4X, 7 ± 1.8%, n = 13, t_(22)_ = 1.5, p = 0.1447; unpaired two-tailed *t* test), suggesting depletion of *Tet1*^*FL*^ or *Tet1*^*S*^ in the dHPC does not affect baseline freezing levels (Fig. 7*F, J*). However, 24 hours later, *Tet1*^*FL*^-SID4X mice exhibited a reduction in their time spent freezing compared to *Tet1*^*FL*^*-*NFD controls (percent time freezing: *Tet1*^*FL*^-NFD, 41 ± 4.7%, n = 12 vs. *Tet1*^*FL*^-SID4X, 26 ± 3.1%, n = 11, t_(21)_ = 2.6, p = 0.0165; unpaired two-tailed *t* test), suggesting impaired memory (Fig. 7*G*). Moreover, *Tet1*^*S*^-SID4X mice exhibited a significant increase in freezing time compared to *Tet1*^*S*^*-*NFD controls (percent time freezing: *Tet1*^*S*^-NFD, 38 ± 2.6%, n = 11 vs. *Tet1*^*S*^-SID4X, 55 ± 3.5%, n = 13, t_(22)_ = 3.8, p = 0.0010; unpaired two-tailed *t* test), indicative of a memory enhancement (Fig. 7*K*). Together, these data demonstrate that hippocampal-dependent memory formation is bi-directionally modulated by the neuron-specific actions of each *Tet1* isoform.

## Discussion

TET1 has been implicated in a wide variety of cognitive functions—most notably learning and memory. However, the exact role of TET1 in the brain has remained ambiguous due to inconsistencies in the findings reported by different studies, suggesting key details regarding its function had yet to be elucidated. In the present study, we report the first definitive evidence that two distinct isoforms of the *Tet1* gene are expressed in the adult mammalian brain. The first, *Tet1*^*FL*^, is transcribed at low basal levels in neurons and encodes for the full-length canonical TET1 enzyme, while the second, *Tet1*^*S*^, is the predominately expressed isoform in the brain, enriched in neurons, and encodes for a recently discovered enzyme variant that lacks a large portion of the N-terminus, including the CXXC DNA binding domain (Zhang et al., 2016). Using isoform-specific genetic tools, we find that the individual disruption of *Tet1*^*FL*^ and *Tet1*^*S*^ has distinct effects on memory formation, excitatory synaptic transmission, and neuronal gene expression, demonstrating that they carry out non-redundant functions.

In the first study to describe *Tet1*^*S*^, it was reported to be the only transcript expressed past early developmental stages (Zhang et al., 2016). Yet, others have recently provided evidence that in some adult tissues and/or cell types, *Tet1*^*FL*^ and *Tet1*^*S*^ are co-expressed (Good et al., 2017; Yosefzon et al., 2017). We found that while *Tet1*^*S*^ is the predominantly expressed isoform in the brain, *Tet1*^*FL*^ is also actively transcribed. We attribute these inconsistencies to differences in the experimental techniques used to detect the lowly expressed *Tet1*^*FL*^ transcript. For example, in the original study, a lack of *Tet1*^*FL*^ expression was inferred using genome-wide RNA-seq data sets, while more recent studies, including our own, directly detected its transcript levels using more sensitive methods such as cell-type specific ChIP-seq, RNAP2 ChIP-qPCR, 5’ RACE and qRT-PCR.

TET1 has been shown to be highly enriched in neurons, but expressed at considerably lower levels in glia (Kaas et al., 2013; Zhang et al., 2013). We find that while *Tet1*^*S*^ is the most abundant transcript expressed in both cell types, in relative terms, the short isoform is expressed at greater levels in neurons (∼3-fold), while in glia, *Tet1*^*FL*^ was ∼15-fold more abundant. Thus, the higher TET1 enrichment previously reported in neurons stems from greater *Tet1*^*S*^ expression, while in glia, despite significantly higher levels of *Tet1*^*FL*^, exhibits lower overall expression due to reduced levels of the shorter transcript. In addition, these findings suggest that the role of each isoform is at least partially cell type-specific. In support of this idea, we found that transcript levels of *Tet1*^*S*^ are downregulated in neurons in response to depolarization and after fear learning, whereas *Tet1*^*FL*^, which is much less abundant in excitable cells, remained steady at baseline levels. Similarly, greater expression of *Tet1*^*FL*^ in glia relative to neurons may reflect its reported role as tumor suppressor in gliomas (Fu et al., 2017), where its added presence is necessary to control gene expression programs related to cellular proliferation.

Several studies using pan-*Tet1* KO mice have found that the loss of both isoforms provides cognitive benefits— including memory enhancement (Rudenko et al., 2013; Kumar et al., 2015; Feng et al., 2017; Cheng et al., 2018). Our findings that acute *Tet1*^*S*^ repression alone is sufficient to enhance long-term fear memory suggests that the pro-cognitive effects observed in previous studies results from loss of the short isoform. In addition, our data are consistent with previous observations that acute overexpression of the TET1 catalytic domain, which also lacks the N-terminal domain of the full length enzyme, impairs memory formation (Kaas et al., 2013). Together, these observations strongly point to *Tet1*^*S*^ as a memory suppressor, and perhaps more generally, as a negative regulator of neuroplasticity. In contrast, acute *Tet1*^*FL*^ repression resulted in memory deficits. In line with our results, a recent study showed that transgenic overexpression of the *Tet1*^*FL*^ gene causes enhanced memory formation and increased anxiety in mice (Kwon et al., 2018), suggesting that overexpression of the full-length enzyme acts oppositely to our SID4X-mediated suppression. It is important to note that while our isoform-specific behavioral findings generally agree with previous reports, several groups have found that pan-*Tet1* deficient mice either have normal memory or exhibit an impairment (Zhang et al., 2013; Towers et al., 2018). Despite our results, the exact cause of these conflicting findings is still not clear. It seems likely, as has been previously posited (for review see (Alaghband et al., 2016; Antunes et al., 2019), that these inconsistencies reflect the use of KO mice with different exons deleted and/or the presence of developmental confounds; the latter being particularly relevant as some *Tet1* mutant alleles display embryonic semi-lethality and/or smaller stature than littermates (Dawlaty et al., 2011; Kang et al., 2015; Towers et al., 2018).

Consistent with our behavioral data, individual disruption of *Tet1*^*FL*^ and *Tet1*^*S*^ in hippocampal neuron cultures had dissimilar effects on excitatory synaptic transmission. In particular, *Tet1*^*S*^ depletion lead to a significant reduction in mEPSC frequency. This presynaptically driven process has been shown to inversely correlate with the strength of long-term depression (LTD) in hippocampal slices (Zhang et al., 2005). Indeed, Rudenko *et al*. reported enhanced LTD in hippocampal slices from pan-*Tet1 KO* mice with strengthened memory retention (Rudenko et al., 2013). LTD has been shown to be necessary and sufficient to facilitate long-term spatial memory formation (Ge et al., 2010; Dong et al., 2012b), thus providing a rationale for how reduced mEPSC frequency might lead to enhanced memory in *Tet1*^*S*^*-*deficient mice. In contrast, acute transcriptional repression of *Tet1*^*FL*^ leads to increased mEPSC frequency and amplitude. While more work is required to resolve why this is, given our behavioral findings after *Tet1*^*FL*^ depletion, this may indicate that *Tet1*^*FL*^ normally acts to suppress aberrant hyperexcitability that leads to impaired cognition.

Our transcriptomic data provides important insights into how *Tet1*^*FL*^ and *Tet1*^*S*^ repression alters neuronal physiology and cognition. For instance, acute loss of *Tet1*^*FL*^ substantially disrupted the expression of a large swath of the neuronal genome, with a significant upregulation of cancer and immune response pathways as well as the down-regulation of genes important for neuronal physiology and learning. These data point to the canonical isoform as a critical regulator of genomic stability in neurons and provides a straightforward explanation for the impaired memory we observed in *Tet1*^*FL*^*-*deficient mice. Likewise, the hyperexcitability in *Tet1*^*FL*^*-*depleted neurons is likely due, in part, to aberrant induction of the immune response. In particular, we found that the expression of genes associated with the Tumor Necrosis Factor (Tnf) pathway to be significantly upregulated. In support of this idea, its activation via the cytokine Tumor Necrosis Factor α (TNFα) has been shown to be sufficient to increase excitability (Ming et al., 2013). Interestingly, previous transcriptomic analyses of constitutive pan-*Tet1* KO have not reported this induction of immune response genes (Rudenko et al., 2013; Zhang et al., 2013; Towers et al., 2018). We propose that the absence of increased inflammation in these studies is likely due to compensatory mechanisms at play during development, as viral-mediated pan-*Tet1* KO in the nucleus accumbens of adult mice has been shown to strongly induce immune gene expression (Feng et al., 2017).

Relative to *Tet1*^*FL*^, *Tet1*^*S*^ inhibition disrupted the expression of far fewer genes. These results are in agreement with data showing that *Tet1*^*S*^ overexpression in HEK293 cells results in significantly fewer DEGs compared to *Tet1*^*FL*^, indicating that it plays a more selective role in gene regulation (Good et al., 2017). The mechanisms behind this difference in gene target specificity remains unclear, but evidence from a recent study of both isoforms suggests it is due, in part, to the absence or presence of the N-terminus, which has been shown to promote global chromatin binding (Zhang et al., 2016). Similarly, given that TET enzymes interact with a large number of DNA-binding proteins, these differences likely reflect their recruitment to specific genomic *loci* by yet-to-be identified co-factors (Good et al., 2017; Wu and Zhang, 2017). In addition, how *Tet1*^*S*^ disruption leads to enhanced memory at the molecular level is not clear. Though, based on our data, it may involve alterations in translation, as loss of the short isoform in neurons leads to a significant up regulation of genes encoding ribosomal RNAs (rRNAs) and proteins involved in ribosomal biogenesis. Consistent with this idea, it was recently reported that learning-induced changes in rRNA expression are required for memory consolidation (Allen et al., 2018). Nevertheless, future studies addressing the molecular functions of *Tet1*^*S*^ and *Tet1*^*FL*^ will be needed to fully understand their roles in regulating nervous system function.

In conclusion, *Tet1*^*FL*^ and *Tet1*^*S*^ are co-expressed in the adult brain, and carry out distinct functions, providing important new insights into the role of TET enzymes in the nervous system. *Tet1*^*S*^ repression enhances memory formation, suggesting that antagonists selective only for the truncated enzyme may be an effective therapeutic strategy to treat cognitive deficits. *Tet1*^*FL*^, on the other hand, appears to be a critical regulator of neuroinflammation and cellular identity, suggestive of a role in aging, neurodegeneration and cancer. Overall, our results stress the importance of distinguishing between the two isoforms in future studies and provide the impetus to reexamine previous findings related to TET1 in depression, addiction and bipolar disorder (Dong et al., 2012a; Feng et al., 2015, 2017).

## Materials and Methods

### Animals

Experiments were performed using 2-3 month old C57BL/6J male mice originally purchased from The Jackson Laboratory. All mice were group housed, kept under 12:12 light/dark cycles, with food and water available *ad libitum*. For stereotaxic surgeries and behavioral assays, 4 week old male littermates were purchased, housed in sets of 3-5 animals and aged out to 10 weeks prior to experimentation in an effort to minimize fighting. All procedures and behavioral assays were approved by the Vanderbilt Animal Care and Use Committee.

### ChIP

Tissue samples were cut into ∼1 mm pieces using a razor and incubated in a 1x PBS solution containing 1% formaldehyde and proteinase inhibitors for 10 min at 37°C, followed by the addition of glycine (final conc. 125 mM) to quench the reaction. Samples were washed 6x with ice-cold PBS and then homogenized in lysis buffer (50 mM Tris pH 8.1, 10 mM EDTA, 1% SDS) using a pestle. Chromatin was sheared using a Bioruptor® Pico set to 3 cycles of 30 s *on*, 30 s *off*. Samples were then processed using the Magna ChIP G Tissue kit (EMD Millipore) according to manufacturer instructions. Briefly, samples were pre-cleared using 25 uL of Protein G beads, then placed on rotator and incubated overnight at 4°C with 25 uL of Protein G beads and 4 uL (1 mg/mL) anti-RNA polymerase II (102660; Active Motif). Immune complexes were sequentially washed according to kit instructions. To reverse cross-links, samples were incubated at 65°C for 2 hrs in the presence of SDS and Proteinase K, then at 95°C for 10 min. Enriched DNA samples were purified with a Qiagen PCR clean up kit. Purified DNA was either stored at -20°C or used immediately for quantitative real time PCR (qRT-PCR). Fold enrichment for each primer set was normalized to input and reported relative to a negative control gene desert region (mouse Igx1a (Qiagen). Primers: *Tet1*^*FL*^ promoter F 5’-gcactctgcaactggtttg-3’, R 5’-gtagaagaggcaggtagaggta-3’; *Tet1*^*S*^ promoter F 5’-ctgctttgaaacaccatgataa-3’, R 5’-tagccatcttgcctgctt-3’.

### 5’ Rapid amplification of cDNA ends (RACE)

Total RNA was extracted from adult mouse hippocampal tissue using an RNeasy Plus Kit (Qiagen). Amplification of 5’ cDNA ends was performed using the GeneRacer™ Kit (Life Technologies) in accordance with manufacturer instructions. Briefly, 3 µg total RNA was dephosphorylated, decapped and then ligated to a GeneRacer™ 5’ RNA oligo. Between each step, samples were purified by phenol:chloroform extraction and precipitated with ethanol and glycogen. Complementary DNA (cDNA) was then synthesized from the ligated RNA using Superscript III™ reverse transcriptase (Invitrogen) and random primers. cDNA samples were amplified for two rounds by nested PCR using Platinum™ PCR Supermix High Fidelity (Invitrogen) and 5’ GeneRacer™ forward and reverse *Tet1-*isoform specific primers (*Tet1*^*FL*^ 5’ RACE outside, 5’-ttgggtgtgactactgggcgctgggaga-3’; *Tet1*^*FL*^ RACE nested, 5’-ggcgctgggagagtcgccagctaaga-3’; *Tet1*^*S*^ 5’ RACE outside, 5’-agccaggcttctggaagagcagggtgt; *Tet1*^*S*^ RACE nested, 5’-cccggaggtggtgacactcatggcatcctt-3’). Purified PCR products were cloned into pCR™-Blunt II-TOPO® vectors (Invitrogen) and sequenced (GenHunter Corp.).

### qRT-PCR

Total RNA was extracted from samples using an RNeasy Plus Mini Kit (Qiagen). RNA samples were eluted in 30-50 µL of RNase free water and analyzed on a NanoDrop™ One Microvolume UV-Vis Spectrophotometer (Thermo Scientific). RNA was then either used immediately for cDNA synthesis or stored at -80°C. cDNA synthesis was carried out in 20 uL reactions using the iScript™ cDNA Synthesis Kit (BioRad) according to manufacturer instructions. All cDNA reactions were diluted 1:5 to reduce potential PCR reaction inhibition with RNase-free water. qRT-PCR was performed on an CFX96 real-time PCR detection system in 10 uL reactions containing SsoAdvanced™ Universal SYBR® Green Supermix and 200-300 uM of primer and 1 uL of cDNA. All qRT-PCR primers were either designed using Primer Quest (Integrated DNA Technologies) to span exon-exon junctions or were acquired directly as pre-designed PrimeTime® qPCR Primer Assays (Integrated DNA Technologies). Relative fold quantification of gene expression between samples was calculated using the delta-delta Ct method and normalized to the geometric mean of the three reference genes *Hprt, Gapdh* and *Gusb*. The comparative Ct method was used to calculate differences in gene expression between samples (Livak and Schmittgen, 2001). IDT PrimeTime qPCR Probe Assays: *Hprt*, Mm.PT.39a.22214828; *Gapdh*, Mm.PT.39a.1

*Gusb* F 5’-cagactcagttgttgtcacct-3’, R 5’-tcaacttcaggttcccagtg-3’;

*Tet1*^*FL*^ F 5’-ctccctggtcatgtacctcta-3’, R 5’-gtaagtaaagatgcaaggatgcg-3’;

*Tet1*^*S*^ F 5’-cctccatctttatttatgcaag-3’, R 5’-ggtttgttgttaaagtctgtct-3’;

*Npas4* F 5’-cacgtcttgatgacaatatgcc-3’, R 5’-ccaagttcaagacagcttcca-3’;

*Arc* F 5’-acgatctggcttcctcattctgct-3’, R 5’-aggttccctcagcatctctgcttt-3’;

*Egr1* F 5’-agcgccttcaatcctcaag-3’, R 5’-tttggctgggataactcgtc-3’.

### Mouse primary neuron, glia and neuroblastoma 2a (N2a) cultures

Mouse hippocampi from C57BL/6J P0 pups were dissected in ice-cold HBSS (Gibco) and digested with papain (Worthington) for 25 min at 37°C. Samples were then washed 3x in HBSS and dissociated by pipetting up and down 15-20 times through a P1000 pipette in growth media (Neurobasal media supplemented with 5 % FBS, 500 nM L-glutamine, B27 (Gibco) and Pen-strep. The cell suspension was passed through a 100 μm filter and centrifuged for 5 min at 500 x *g*. Cell pellets were resuspended in growth media and seeded on poly-d-lysine coated (Sigma) 12-well plates (Corning) at ∼250-300 × 10^3^ cells per well or on 24 well plates at 100 × 10^3^ cells per well. 24 hours later, media was replaced with maintenance media (Neurobasal media supplemented with 500 nM L-glutamine, B27) without Pen-strep. At DIV 3-4 1 µM 5-fluorodeoxyuridine (FdU) was added to the media for 24 hours, then removed, to inhibit mitotic cell growth (Hui et al., 2016). Pharmacological stimulation of primary hippocampal neurons was carried out on DIV10-12. Briefly, 25 µL of maintenance media alone (vehicle), containing KCl, Bicuculline or NMDA/glycine (final conc. 25 mM, 50 µM, 10 µM/2 µM, respectively) were administered to each well and incubated for 1, 2, or 4 hrs. Wells were then washed with 1x PBS, processed immediately or stored at -80°C. Primary mixed glial cultures were prepared identically to neurons, except they were plated and maintained using media that consisted of 1x DMEM (+4.5g/L D-glucose and L-Glutamine, Gibco), Pen-strep and 10% FBS. DIV14 confluent glial cultures were used for qRT-PCR experiments. Neuro-2a cells (N2a) were purchased from ATCC, and cultured using the same media as glial cells. For transfection experiments, N2as were seeded in 24 well plates at a density of 200 × 10^3^ per well and transfected using GenJet™ Reagent (II) in accordance with manufacturer instructions.

### Vector construction

All constructs used in this study were generated using Gibson Assembly methodology as previously described (Gibson et al., 2009). Briefly, primers were designed using NEBuilder Assembly Tool (v1.12.18) to generate overlapping (20-25 bps) PCR fragments amplified using Q5® High-Fidelity DNA Polymerase (NEB). Modified TALE constructs were created using PCR fragments amplified from pAAV-CW3SL-EGFP (Addgene # 61463), a gift from Bong-Kiun Kaang (Choi et al., 2014), and constructs contained in the TALE Toolbox (Addgene kit # 1000000019), a gift from Feng Zhang. TALE DNA targeting sequences were assembled into TALE construct backbones using Golden Gate Assembly Cloning, as previously described (Sanjana et al., 2012). TALE sequences targeting the *Tet1*^*FL*^ (5’-TGCCCCAGCTACACTCCT-3’, sense) and *Tet1*^*S*^ (5’-TCGCAGCCTAGCACTATC-3’, antisense) promoter regions were designed using TAL Effector Nucleotide Targeter 2.0 software (Doyle et al., 2012).

### Immunostaining

For immunocytochemistry, cells were plated on glass cover slips coated with poly-d-lysine. Cells were washed 2x with ice cold 1x PBS, followed by fixation with fresh 4% PFA for 15 min at room temperature, blocked for 1 hr at room temperature (10% goat serum + 0.3% Triton-X 100), and incubated with primary antibodies (anti-HA, ab18181, abcam; anti-GFP, ab13970, abcam; anti-NeuN, ab104224) at a concentration of 1:1000 at RT for 2 hr or overnight at 4°C in 1:3 diluted blocking buffer. Slides were then washed and incubated with the appropriate Alexa flour secondary antibodies (Abcam) at a concentration of 1:1000 for 1 hr at RT. Slides were mounted using ProLong™ Gold Antifade Mountant with DAPI (Invitrogen) and images were acquired using a IX73 microscope (Olympus) and cellSens standard software.

### AAV generation and viral injections

High titers (>10^12^ gc/mL) of AAV1 viral particles containing *Tet1* isoform TALE constructs were packaged by Applied Biological Materials (ABM). For primary hippocampal neuron experiments in 12 or 24 well plates, we added 1 uL of AAVs diluted 1:10. For *in vivo* experiments involving stereotaxic surgeries, AAVs were injected bilaterally into the dorsal hippocampus of 10 week old mice using the following stereotaxic coordinates: –2.0 mm AP, ±1.5 mm ML, and -1.6 mm DV from bregma. A total of 1.5 μL of viral solution was injected per hemisphere. Injections were performed using a 10 mL Hamilton Gastight syringe controlled by a Pump 11 Elite Nanomite Programmable Syringe Pump (Harvard Apparatus). Injections proceeded at a speed of 150 nL min^−1^ through a 32-gauge needle. The injection needle was left in place an additional 5 min. Behavioral or gene expression experiments involving AAV delivery *in vivo*, were either begun or performed 14 days post-surgery, respectively.

### Behavior

All behavioral experiments were carried out within the Vanderbilt Mouse Neurobehavioral Core facilities (https://lab.vanderbilt.edu/mouse-core/). After all behavioral experiments were completed, the brains of all animals were removed for tissue collection and analysis of AAV infection localization within CA1-CA3 hippocampal subfields under an IX73 microscope (Olympus). Due to the reported role of TET1 in neurogenesis, any animals that exhibited AAV-mediated EGFP expression in the dentate gyrus were excluded from data sets to remove potential confounds (Zhang et al., 2013).

#### Elevated Zero Maze (EZM)

Mice were placed on the open section of the maze (White 2325-0229, San Diego Instruments) and allowed to explore freely for 5 min. Video recording and tracking were performed using ANY-maze video tracking software (Stoelting Co.).

#### Open field

Mice were placed in the center of a large Plexiglas box (43 × 42 × 30 cm) and locomotor activity was measured for 30 min (Med Associates, Inc.). Data is presented as total distance traveled in centimeters.

#### Contextual Fear conditioning (CxFC)

Fear conditioned mice used for behavioral analysis were trained in a novel context (Cat# MED-VFC2-SCT-M, Med Associates Inc.) using a 3.5 min training protocol consisting of a 3 min habituation period, followed by a single foot shock (0.5mA, 2 s). Mice were removed from the chamber 30 s later. To assess long-term memory formation, mice were placed back in the same context for 5 min in the absence of the unconditioned stimulus. Percent freezing was calculated automatically using Video Freeze Version 2.1.0 (Med Associates Inc.). Fear conditioned animals used for gene expression analysis were trained using a protocol consisting of three context-shock pairings as described above (0.75mA, 2 s) every 2 min, and removed from the apparatus after 7 min.

### RNA-seq and ChIP-seq

For RNA-seq, total RNA was extracted from AAV1-transduced primary hippocampal neurons (DIV12-14) using an AllPrep DNA/RNA/Protein Mini Kit (Qiagen). Total RNA was polyA selected and sequenced (Hudson Alpha GSL) on the Illumina platform (HiSeq v.4, paired end, 50 bp, 50 million reads). Reads were passed through a quality filter with trimmomatic (Bolger et al., 2014) using recommended settings for paired end libraries, and adaptor sequences matching TruSeq3_PE were trimmed. Surviving reads were aligned to the mm10 genome with hisat2 (Kim et al., 2015). Stringtie (Pertea et al., 2015) was used to incorporate any novel transcripts from these sequencing libraries into the mm10 Refseq annotation of known transcripts, and featurecounts (Liao et al., 2014) attributed reads to the resulting custom annotation. EdgeR (Robinson et al., 2009; McCarthy et al., 2012) was used for determining fold change and false discovery rates (FDR) for each gene with sufficient read depth. Statistics in EdgeR were determined with genewise negative binomial generalized linear models with quasi-likelihood tests (glmQLFit function). H3K4me3 ChIP-seq from CA1 neurons was downloaded from (Halder et al., 2015). Alignment, peak calling, and file visualization were conducted as described in Collins et al., 2019. RNA-seq datasets generated in this study have been deposited in Gene Expression Omnibus (GEO) with the accession number GSE140174.

### Electrophysiology

Whole-cell voltage clamp recordings were performed on neurons from 14 - 21 DIV mouse hippocampal cultures using a Multi-Clamp 700B amplifier. Signals were digitized through a Digidata 1440A at 20 kHz, filtered at 1.8 kHz, and analyzed offline with Clampfit 10.7 software (Molecular Devices). Cells were held at -60 mV. Patch pipettes were pulled from borosilicate glass capillaries with resistances ranging from 3 - 6 MΩ when filled with pipette solution, containing (in mM/L):120 Cesium Methanesulfonate, 5 CsCl, 1 MgCl_2_, 1 CaCl_2_, 10 HEPES, 2 ethylene glycol-bis-(aminoethyl ethane)-N,N,N’,N’-tetraacetic acid (EGTA), 4 Na-ATP, 0.4 Na-GTP, 10 phosphocreatine, 3 Na-Ascorbate, 5 Glucose, pH 7.2. The bath solution (Tyrode’s saline) contained (in mM/L):150 NaCl, 4 KCl, 2 MgCl_2_, 2 CaCl_2_, 10 N-2 hydroxyethyl piperazine-n-2 ethanesulphonic acid (HEPES), 10 glucose, pH 7.35. For recordings of miniature excitatory post-synaptic currents (mEPSCs), bath solution was supplied with1 µM tetrodotoxin (TTX, Hello Bio). mEPSC events were collected from the recorded traces using a template-based approach. Templates was generated from the trace recorded in control conditions. 30 randomly selected events from individual recordings were used for analysis.

### Experimental Design and Statistical Analyses

Statistical analysis and graphing of non-genomic data was performed using Graphpad prism 7.04. For two groups, statistical significance was determined using the Student’s *t* test. For 3 or more groups, statistical significance was determined using One or Two-way ANOVA, followed by Dunnett’s or Sidak’s multiple comparisons tests, respectively, *post hoc*.

## Supporting information

Extended data Tables related to figure 5

## Author contributions

The study was conceived by G.A.K. Experiments were coordinated with help from J.D.S., and A.J.K. Vectors were generated and tested by G.A.K with help from J.Z. and K.S.C. Molecular experiments, stereotaxic surgeries and behavioral tests were carried out by G.A.K. with help from, J.W. J.D.W. and A.Y.J. Ephys experiments and analysis was performed by S.P.M and R.M.L. RNA-seq experiments and data analysis were performed by C.B.G. The paper was written by G.A.K and C.B.G, with help and comments from all other authors.

## Acknowledgements

Funding support for this study was provided by NIH grants MH057014 and MH107254 to J.D.S., P20-GM103423 to A.J.K., T32-MH065215, 1S10ODO17997 to the VMSRC, and U54-HD083211 to the VMNL). We thank Hehuang Xie, Roger J. Colbran, Danny G. Winder and Colleen Niswender for helpful comments and discussions during the preparation of this manuscript. Finally, we would like to acknowledge the Murine Neurobehavioral Core lab at the Vanderbilt University Medical Center for their guidance and use of equipment.

## Extended Data

**Extended data table 5-1. All DEGs between AAV1-*Tet1*^*FL*^-SID4X versus AAV1-*Tet1*^*FL*^-NFD**. Significantly changed genes by repression of *Tet1* ^*FL*^. Column headings: MSTRG – Stringtie ID or Refseq ID of gene (if gene prediction matches annotation exactly); logFC = log2 fold-change; logCPM = average log2 counts per million; F – F statistic; PValue = probability value before multiple hypothesis correction; FDR = false discovery rate; Refseq = best matching Refseq gene ID that best matches the stringtie predicted transcript; OGS – official gene symbol with period after to prevent date conversion in excel; Sig – whether the transcript passes the FDR and fold change cutoffs for significance.

**Extended data table 5-2. DEGs significantly activated by AAV1-*Tet1*^*FL*^-SID4X relative to AAV1-*Tet1*^*FL*^-NFD**. The subset of genes that are activated by repression of *Tet1* ^*FL*^. To the right, the significantly enriched (Benjamini-Hochberg adjusted p values < 0.05) DAVID GO enriched terms for biological process (top) and KEGG pathways (bottom). The top ten terms included in the main figure are bolded. Column headings: Category – gene ontology set utilized; Term – gene ontology term title; Count – number of genes in gene set present in the genes that comprise the ontology term; % -percent of genes in the list of genes for a gene ontology term; P-Value – probability value; Bonferroni – Bonferroni corrected p value; Benjamini – Benjamini-Hochberg corrected p value.

**Extended data table 5-3. DEGs significantly repressed by AAV1-*Tet1*^*FL*^-SID4X relative to AAV1-*Tet1*^*FL*^-NFD**. The subset of genes that are repressed by repression of *Tet1* ^*FL*^.

**Extended data table 5-4. All DEGs between AAV1-*Tet1*^*S*^-SID4X versus AAV1-*Tet1*^*S*^-NFD**. Significantly changed genes by repression of *Tet1* ^*S*^.

**Extended data table 5-5. DEGs significantly activated by AAV1-*Tet1*^*S*^-SID4X relative to AAV1-*Tet1*^*S*^-NFD**. The subset of genes that are activated by repression of *Tet1* ^*S*^.

**Extended data table 5-6. DEGs significantly repressed by AAV1-*Tet1*^*S*^-SID4X relative to AAV1-*Tet1*^*S*^-NFD**. The subset of genes that are repressed by repression of *Tet1* ^*S*^.

**Extended data table 5-7. DEG relative to each NFD control overlap**. The genes that make up the total gene counts in the Venn diagrams in panels C and D.

**Extended data table 5-8. All DEGs between AAV1-*Tet1*^*S*^-SID4X versus AAV1-*Tet1*^*FL*^-SID4X**. Significantly changed genes when repression of *Tet1* ^*S*^ is directly compared to repression of *Tet1*^*FL*^.

**Extended data table 5-9. DEGs significantly repressed by AAV1-*Tet1*^*S*^-SID4X relative to AAV1-*Tet1*^*FL*^-SID4X**. Significantly suppressed genes when repression of *Tet1* ^*S*^ is directly compared to repression of *Tet1*^*FL*^.

**Extended data table 5-10. DEGs significantly activated by AAV1-*Tet1*^*S*^-SID4X relative to AAV1-*Tet1*^*FL*^-SID4X**. Significantly activated genes when repression of *Tet1* ^*S*^ is directly compared to repression of *Tet1*^*FL*^.

